# Fish locomotor variation: connecting energetics and kinematic modulation

**DOI:** 10.1101/2025.07.03.663096

**Authors:** Yangfan Zhang, Divya Ramesh, Hungtang Ko, George V. Lauder

## Abstract

Analyses of vertebrate locomotion have frequently revealed variations in locomotor energetics and movement both among individuals and through time within an individual. This variation is often collapsed into mean values for broad comparative analyses of function. However, kinematic patterns of locomotion, even when animals move at a near-constant mean speed, frequently vary with both the physical and biological context. Here we demonstrate, using analyses of fish locomotion and energetics, how variation among individuals in kinematic gaits can manifest as changes in dynamics of metabolic rate (estimated from oxygen uptake). We present kinematic data from a small school of giant danio (*Devario aequipinnatus*) to show that fish within a school frequently modulate their kinematics and change position, even when the school moves at an overall constant mean speed. We show that rainbow trout (*Oncorhynchus mykiss*), swimming over a range of speeds, exhibit considerable variation in tail beat frequency and metabolic rate among speeds. By experimentally altering the fluid dynamic environment, we demonstrate that brook trout (*Salvelinus fontinalis*) show correlated modulation of both kinematics and metabolic rate. Simultaneous measurement of energetic and biomechanical characteristics can unveil the physiological, biomechanical, and fluid dynamic mechanisms that underlie dynamic changes in vertebrate locomotor gaits.

## 1. Introduction

Biological variation is fundamental to understanding biodiversity. Diverse structures and morphologies accommodate biological functions that can be key for animals’ lifetime fitness. One of the most evident features of animal diversity is variation in movement ability across species [1] [2]. Locomotion within an individual is also dynamic: animals modulate their patterns of movement as they respond to various external stimuli. Canonical examples of locomotor dynamics include roller coaster migration over the Himalayas by bar-headed geese [3], speed-dependent locomotion gaits and their distinct metabolic costs in horses [4], acceleration-and-gliding in diving mammals [5] [6], intermittent locomotion in sharks [7], and burst-and-coast in perciform fishes [8] [9]. These studies directly demonstrate that animals can use multiple locomotion gaits and engage in unsteady-state locomotion [10] [11] [12], which is contrary to the typical presentation of steady-state locomotion in the majority of the literature. Yet, we currently have a limited understanding of why animals exhibit variation and what the metabolic effects of intra-individual kinematic variation in locomotion are. Are variations in locomotor dynamics biologically meaningful, and do they connect to changes in metabolic energy use?

In this regard, it is important to understand the physics and physiological systems underpinning complex transitions among locomotion gaits when animals respond to environmental cues that can potentially achieve metabolic energy saving. Thus, studies of locomotor dynamics focus on how forces act on the motion of bodies and how they change the movement trajectory of a body, coupled with the understanding of metabolic cost.

The integration of biomechanical and bioenergetic data has yielded fruitful results in studies of aerial and terrestrial locomotion in animals [13] [14] [15] [16] [17] and human exercise physiology [18] [19], whereas a conceptual connection to studies of aquatic locomotion has long been considered at the ecological level [20] [21] [22] [23] [24]. In general, few studies simultaneously and synchronously measured the biomechanics and the energetics, regardless of aerial, terrestrial and aquatic locomotion, and despite the importance of integrating biomechanics and energetics, which has recently been highlighted in a special issue of *J. Exp. Biol*. [25].

One key challenge has been the temporal precision of measuring whole-animal metabolic rate and mapping the often rapid metabolic changes to specific locomotor gaits. Whole-animal metabolic rate measurements provide a direct estimate of the metabolic costs of locomotion, moving beyond measurement of muscle power in a few locomotor muscles, or other indirect metrics to allow qualitative comparisons of metabolic costs. Animals have many other tissues using ATP during locomotion, including the cardiorespiratory system and other peripheral muscle groups for postural control, and to determine the overall metabolic cost, it is important to assess whole-organism tissue energy use. Kinematic data, ideally measured with multiple cameras to provide spatial resolution, are of special importance when metabolic measurements are made, as fish are adept at finding zones of beneficial flow and altering their metabolic rate [26]. Moreover, resolving locomotor dynamics needs longer duration high-speed videography for transitions among multiple locomotion gaits. These challenges are currently being mitigated by new experimental methodologies, improved analytical approaches, and enhanced technologies that allow the measurements of aerobic metabolic rate at a higher resolution [27] [28] and estimation of the total energy expenditure of locomotion [29] [26] [30], while at the same time tracking fish kinematics in relation to hydrodynamic flow patterns.

Herein, we focus on illustrating the importance of variation in patterns of fish locomotion and how variation or modulation of locomotor kinematics is reflected in changes in metabolic rates. We provide new experimental data from both fish in schools as well as individual fish swimming at different speeds and under different hydrodynamic conditions. The overarching question is whether modulated locomotor dynamics can enable a reduction in metabolic energy costs during locomotion, even for short periods? Our goal is to contribute to the understanding of the effects of fish locomotor variation at the levels of both the individual and collective, and illustrate an integrated approach to studying the kinematics, energetics, and fluid dynamics of fish movement.

## 2. Research approach

The case studies presented here encompass fish locomotor variation both at the level of an individual and in a collective. The first case study aims to understand fish locomotor variation within an animal collective. The second case study demonstrates tightly coupled biomechanics and energetics of a fish when responding to different fluid flow fields. The methods below provide a general overview of this approach, and details of experimental methods and analytical approaches are provided in the supplementary materials.

### (a) Kinematics

An aquatic vertebrate swim tunnel was used to provide laminar flow (*U* = ∼3 BL s^-1^) to stimulate the directional collective movement of a fish school (8 individuals within a school, ∼6 cm individual total length). To mitigate fish seeking fluid refugia in the boundary layer close to the walls of the swim tunnel, we customized a rectangular enclosure (30 • 11.5 • 11 cm) made with a carbon fibre frame and attached nylon netting [31]. The enclosure was situated at the centre of the testing section (50 • 30 • 30 cm) of the swim tunnel. The ventral view was recorded using one camera at 60 frames per second for ∼3.5 hours (AOS Promon U1000, AOS Technologies, Baden, Switzerland) to continuously record variation in kinematics of individual fish within the school (Fig. 1). Prior to the experiment, the fish school was left undisturbed in the swim tunnel overnight to habituate to the testing chamber. Analyses of individual locomotor kinematics used a published computer vision analysis pipeline [31] (*see* supplementary for detailed methods). This analytical pipeline enables the extraction of individual fish tail beat amplitude (Amp_tail_) and frequency as well as the midline kinematics when the fish are not overlapping in the camera view.

**Figure 1.**
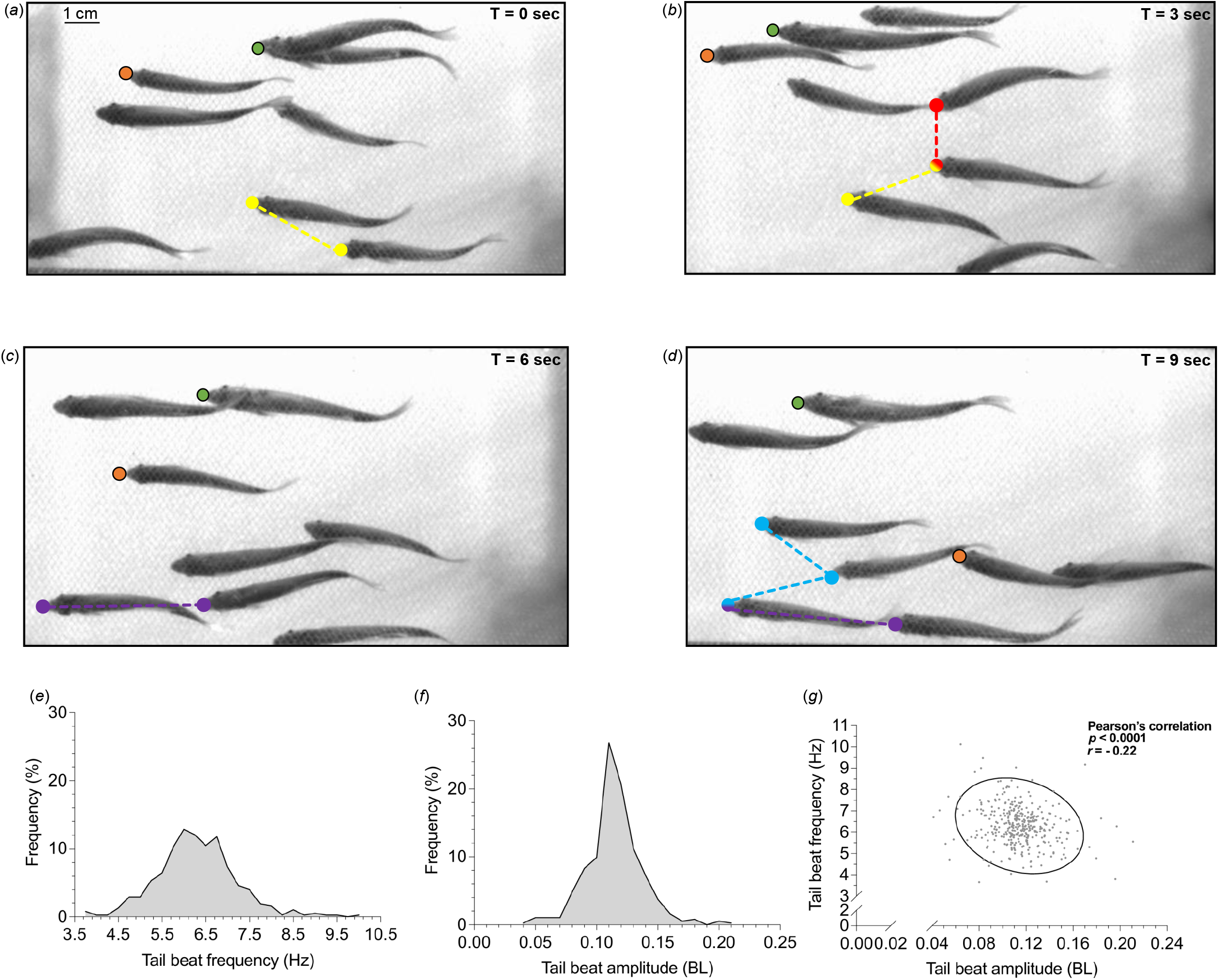
Variation among individual fish within a school in locomotor kinematics and position. (a-d) We qualitatively demonstrate the variation in fish positions using images from video recordings (*see* supplementary video) of a fish school (giant danio, *Devario aequipinnatus*) over 9 seconds (*see* time stamps on the top right of each panel). The orange and green dots (with a solid black border) mark the same two fish across the frames. These two fish showed variation in kinematics and position within the school. The dots (without border) and connected by dashed lines denote fish in various relative positions within the school that have been theoretically proposed or experimentally proven to have hydrodynamic benefits that reduce locomotor energetic cost. Yellow dots indicate fish in a staggered formation; red dots indicate fish side-by-side; purple dots indicate fish in-line. Light blue lines show fish in a triangle formation. One individual fish can be a part of two formations (*see* mixed colour dots). Moreover, we quantitatively demonstrate the variation of locomotor kinematics. (e, f) Frequency distribution of tail beat frequency and amplitude (body length, BL) measured using a contrast-based segmentation algorithm from individual danio within a school (215 minutes video recorded at 60 fps, water velocity ∼3 BL s^-1^). (g) Scatter plot of tail beat amplitude and frequency with a 95% confidence ellipse.

### (b) Energetics

To integrate the analysis of the biomechanics and energetics of a fish, we used an established Integrated Biomechanics and Bioenergetic Assessment System (IBAS) [30] (*see* supplementary material). This system simultaneously measured time-synchronized high-speed videography with dynamic changes in aerobic metabolic rate. Aerobic metabolic rate was estimated from the rate of oxygen consumption by a trout in a sealed respirometer using data obtained from a high-resolution fibre optic O_2_ probe. After habituating a trout to the testing arena overnight, water velocity was gradually increased to 50% of the maximum sustained swimming speed (*U*_crit_) of brook trout (*Salvelinus fontinalis;* mass*=*267g; length*=*26 cm) [31]. Water velocity was maintained at the 50% *U*_crit_ for ∼7 minutes before modifying the fluid flow field (*see* supplementary for detailed methods). To test the effect of altered flow fields on fish swimming, we used a rotating vertical airfoil sealed inside the respirometer while keeping free-stream velocity the same. This airfoil, oriented at an angle of 30 degrees to the oncoming flow, produced a strong velocity gradient between the region behind the foil and the free stream region in the respirometer. The trout volitionally entered the region of the high-velocity gradient and was able to hold a position within the gradient using altered body kinematics compared to free-stream locomotion using undulatory body movement. We continued the kinematic and energetic measurements for ∼12 minutes while fish was in fluid velocity gradient.

## 3. Inter-individual variation in kinematics within a fish school

Fish swimming within schools routinely change relative positions with other individuals [30] [32] [33] [34] and display considerable variation in body kinematics [33] [35]. What remains perplexing is that despite the dynamic changes in the relative positions of individual fish within schools, fish schools as a whole can still achieve energy savings compared to individual fish swimming alone [36] [30] [31]. Fish within a school exhibit positional modulation (Fig. 1a-d) that includes side-by-side positions, staggered configurations, ladder formation [33], as well as in-line tandem positions with one fish behind another [37] [38]. Both biomechanical studies and computational fluid dynamics simulations suggest that fish in each of these positions can reduce their metabolic energy consumption. As demonstrated previously [28], fish swimming in a school display reduced the cost of locomotion as a collective, especially at higher speeds, relative to an individual swimming alone.

One noteworthy aspect of fish schooling dynamics is the considerable variation that occurs in body kinematics by fish within the school. Although previous research has identified changes in tail beat frequency (*f*_tail_) by fish located at the back of a school compared to fish near the front [39], for the most part, fish within a school are treated as exhibiting generally similar undulatory body kinematics, which might not be the case. Using computer-vision-based tracking techniques [32] [33] [40] [41], we analyzed data from long-term video recordings of fish swimming in a school (Figs. 1 & 2) to characterize the extent of kinematic variation among individual fish and the extent of modulated locomotion. Long-term continuous kinematic data on fish schooling behaviour remain scarce and sharply contrast with the typically short time scale (seconds to minutes) of most kinematic data on fish behaviour.

Individual fish within the school show kinematic modulation during nearly four-hour-long recordings of the kinematics of individual fish within an 8-fish school (at 60 frames per second). Although the average *f*_tail_ was 6.3 Hz, *f*_tail_ ranged from 3.7-10.1 Hz with 25-75 percentile of 5.8-6.8 Hz and a coefficient of variation of 14.5%. Only 11.5% of the time, individual fish within a school exhibited the average *f*_tail_ of 6.3 Hz (Fig. 1e). Likewise, 27% of the time, individual fish within a school exhibited the average Amp_tail_ of 0.11 BL, showing a range of Amp_tail_ from 0.041 to 0.17 BL with a 25-75 percentile of 0.1-0.13 BL and a coefficient of variation of 19.7% (Fig. 1f). Less than a quarter of the time, fish within the school were swimming with the kinematic pattern of group average. Thus, considerable intra-and interindividual variation in *f*_tail_ and Amp_tail_ is present within a school. Moreover, individual fish within a school at various times can reduce their *f*_tail_ and Amp_tail_ to levels that are substantially lower than the group average. If a lower *f*_tail_ reflects a reduced energy expenditure (despite certain caveats [26] [30]), we hypothesize that individual fish within the school appear to momentarily reduce their locomotor energy consumption by occupying positions that require less energy, while still holding position in the streamwise direction. A reduction in required energetic expenditure could also result from fish moving from the front of the school toward the back (Fig. 1, fish identified by the orange dot) by reducing *f*_tail_ and/or Amp_tail_ to generate less thrust and allowing body drag to cause a change in position within the school.

Fish within the school also varied considerably in Amp_tail_, and individuals with a high *f*_tail_ could display reduced midline oscillation and a lower Amp_tail_ (Fig. 2b, f), while other individuals with high *f*_tail_ showed increases in Amp_tail_ (Fig. 2c, g). Fish with a lower *f*_tail_ (Fig. 2d, e) could also show both increased and reduced Amp_tail_ (Fig. 2h, i), as well as asymmetrical kinematics when compared to individuals swimming relatively steadily and not modulating body position (Fig. 2e). These examples demonstrate that even when fish use a similar *f*_tail_, midline kinematics can vary substantially, and that modulation of Amp_tail_, *f*_tail_, and the undulatory wave all occur within a school. There is a possible negative trade-off between *f*_tail_ and Amp_tail_ within the school (Fig.1g; Pearson’s correlation coefficient: *p* < 0.0001 *r* = -0.22). Currently, long-duration and simultaneous measurements of *f*_tail_ and Amp_tail_ are few in the literature and hinder our capacity to draw comparisons. Hence, more studies are needed to further understand whether or not a limit exists for the individual fish within a fish school to increase both *f*_tail &_ Amp_tail_. Although the burst swimming of a solitary fish shows both increased *f*_tail_ and Amp_tail_ [42], steady swimming typically involves increases in *f*_tail_ while maintaining Amp_tail_ to achieve energetic efficiency [43] [32] [44]. A systematic understanding of the potential negative trade-offs between *f*_tail_ and Amp_tail_ is lacking, although the trade-off of *f*_tail &_ Amp_tail_ that we observed might relate to the limit on the contracting and bending properties of the musculature as well as the positional modulation of individual fish within a school.

**Figure 2.**
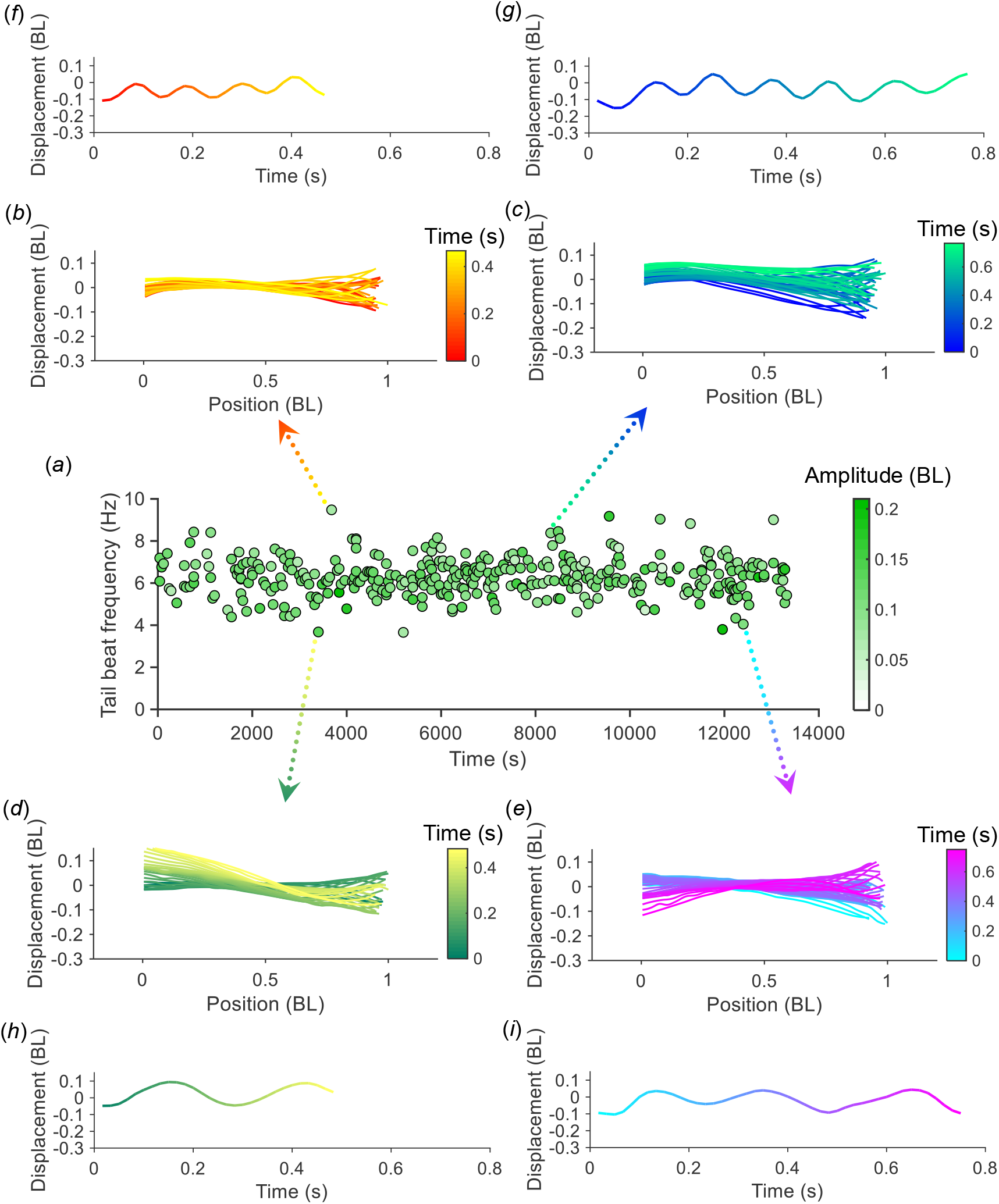
Variation in fish kinematics within a school (giant danio, *Devario aequipinnatus*) was recorded continuously over a time of more than 215 minutes. (a) Tail beat frequency of individual fish with a fish school (each point is one individual with a 40-sec bin). Thus, individuals are measured repeatably in each subsequent 40-sec bin. The regression model estimated the average tail beat frequency as 6.3 Hz (at ∼3 BL s^-1^), whereas inter-individual variation was considerable, with a range of 3.7-10.1 Hz. The colour of each data point corresponds to the peak-to-peak amplitude of the tail tip or base, depending on the segmentation of the caudal fin. (b) Midline kinematics of an individual fish with a high tail beat frequency (9.5 Hz) (duration: 0.47 sec, number of midlines: 28) closely matching mean flow speed and not changing position within the test chamber; (c) Midline kinematics of an individual fish with a high tail beat frequency showing lateral movement (8.4 Hz) (duration: 0.77 sec, number of midlines: 46); (d) Midline kinematics of an individual fish with a low tail beat frequency and with large lateral body oscillation displacement (3.7 Hz) (duration: 0.48 sec, number of midlines: 29); (e) Midline kinematics of an individual fish with a low tail beat frequency (4 Hz) while maintaining a stationary position in the test chamber (duration: 0.75s, number of midlines: 45). (*f-i*) The oscillatory pattern of tail displacement, as a function of time, is colour-matched to the corresponding midline kinematics with a peak-to-peak tail beat amplitude of (*f*) 0.08 body length (BL), (*g*) 0.1 BL (*h*) 0.14 BL, and (*i*) 0.11 BL.

## 4. Variation in kinematics and energetics with speed

Biologically meaningful variation is not only present at the level of the animal collective but also manifests as modulated locomotion at the individual level. This variation can present both across movement speeds as well as within a given speed.

The concept of locomotor performance curve [26] [34] describes the changes in the performance metrics of interest (such as Amp_tail_, *f*_tail_, and energy use) as a function of speed. Despite dynamic changes in the locomotor kinematics, metabolic locomotor performance curves are often presented as the average animal metabolic rate at each speed, even though there is variation both in metabolism and kinematics at each speed. A key challenge is to temporally match changes in metabolism with the measurements of locomotor mechanics at a fine scale to permit the analysis of the relationship between kinematics and metabolic energy at each speed. Metabolic rate is often presented as average values because whole-animal metabolic rate measurements have traditionally needed almost 10 mins to stabilize at a reliable value. Each step of the established critical swimming speed protocol (*U*_crit_) [29] can require at least 10 minutes [29] [31] [45].

As the measurement techniques of whole-animal metabolic rate advance in resolution, the metabolic costs quantified by the locomotor performance curve can be viewed at a higher fidelity. In particular, a modern optical oxygen probe can reduce the required measurement window to nearly 1.5 minutes and has a measurement frequency of 1 Hz [27]. Using a sliding window algorithm, whole-animal metabolic rate can be measured at the resolution of one measurement per second, which enables a measurement of dynamic metabolic rate [28]. Data obtained from rainbow trout using the dynamic metabolic rate approach [27] shows that *f*_tail_ is variable (coefficient of variation: 7.8-12.1%; Fig. 3a: grey dots) even within the same flow velocity. The metabolic rate of the fish, correspondingly, is also dynamic as shown either by the average (black points) or as the variation (grey dots) (Fig 3b). Therefore, within each movement speed of the locomotor performance curve, biological variations in both kinematics and energetics occur and are correlated: fish that show an increase in *f*_tail_ also demonstrate increased metabolic cost.

**Figure 3.**
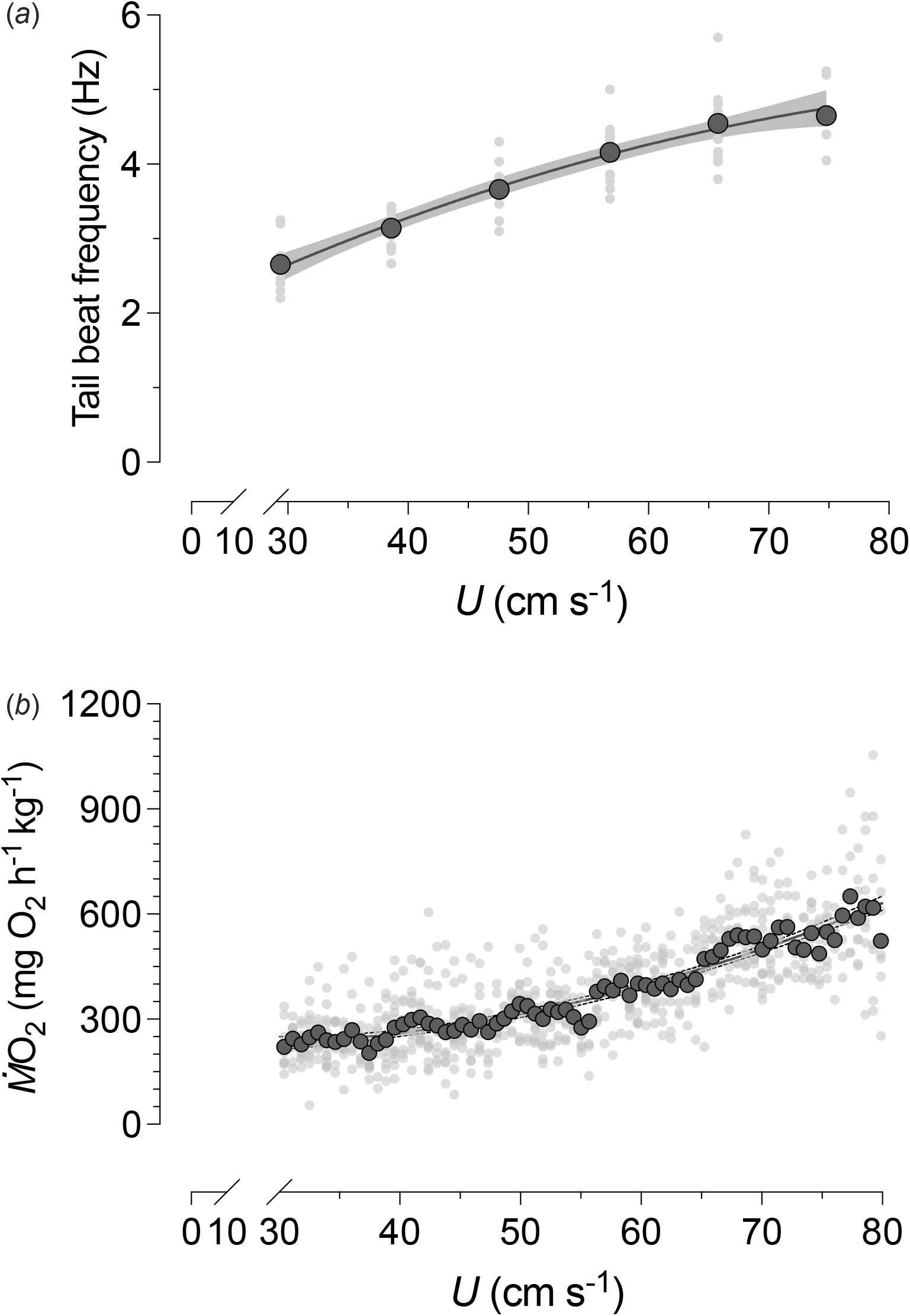
Kinematics and dynamic metabolic rate of a fish in a critical swimming speed (*U*_crit_) testing protocol were measured simultaneously to illustrate the extent of variation in both variables. (a) The tail beat frequency of rainbow trout (*Oncorhynchus mykiss*) (∼70g) during a *U*_crit_ protocol in laminar flow. In addition to the average (point with solid lines), the tail beat frequency of an individual fish showed a range of tail beat frequencies (grey dots) at given speeds. The relationship between the average tail beat frequency and the speed is presented as a solid curve (y = 0.198 + 0.096x – 0.00047x^2^; R^2^ = 0.79, AIC = -142.5) with a 95% confidence interval (shaded area). (b) The aerobic metabolic rate is dynamic across and within each flow speed. Besides the average dynamic aerobic metabolic rate (dots with a solid black border), each individual (grey dots without border) also showed variation. The relationship between average dynamic aerobic metabolic rate and speed is represented by a solid curve (y = 299.2 – 6.1x + 0.13x^2^; R^2^ = 0.61, AIC = 6852) with a shaded 95% confidence interval (dashed lines). The data were adapted from [27].

Burst-and-coast swimming is a well-studied gait that represents an important case of modulated locomotion in fish. The occurrence of burst-and-coast locomotion increases when fish are approaching and exceeding a gait transition speed that is typically 50% of the maximum sustained swimming speed [8] [9]. Burst swimming is powered by fast-twitch white muscle fibres that predominantly use the ATP generated by anaerobic glycolysis [29] [9] [8] [46] [47].

The coasting phase, which shows close to zero *f*_tail_, is shown to help replenish the venous oxygen content [48]. Burst swimming gaits correspond to peaks in the aerobic metabolic rate, whereas the coasting phase corresponds to the troughs in the aerobic metabolic rate [27]. Although more research is needed, one current hypothesis is that the bursting phase provides the power needed to counter fluid drag as swimming velocity increases. Indeed, biomechanical and hydrodynamic analyses suggest that fish tail acceleration during the bursting phase can boost the efficiency of propulsion by altering the geometry of the vortex ring to increase the velocity-power ratio [42] [49]. The coasting phase provides brief moments to recover high-energy phosphate substrates and replenish the oxygen stores in blood haemoglobin and muscle myoglobin [50]. Therefore, the burst-and-coast swimming exemplifies the deep connection between cardiorespiratory physiology and metabolism with biomechanics in animal locomotion.

## 5. Locomotor variation resulting from dynamic fluid conditions

The majority of controlled laboratory studies in fish locomotion expose fish to laminar flow, and strive to limit changes in flow conditions to fluid velocity alone, even though it is widely recognized that fluid movement in nature is far more complex [51] [52] [53] [54]. Flow regimes can be affected by solid obstacles that create high-drag regions behind objects as well as by fluid separation and velocity gradients that occur at the edges of obstacles. Animal locomotion in nature needs to navigate around, cope, and even take advantage of such fluid velocity gradients, and much of the locomotor variation that we observe in fish can be a response to altered fluid dynamic conditions [55]. Variation in fluid dynamics is thus responsible for inducing changes in fish kinematics and metabolism.

One classic example of fish dealing with obstacles in flow that alter fluid dynamics and affect body kinematics and metabolism is the change that occurs when fish interact with the drag wake (or Kármán wake) behind an obstacle [56]. When fish interact with a Kármán vortex street (a rhythmical oscillation in velocity resulting from vortex shedding behind a blunt obstacle), they show altered body kinematics, lower muscle activity levels [56], and reduced metabolic costs of locomotion [57].

Fish also frequently encounter velocity gradients and must deal with accelerated flows. Velocity gradients create pressure gradients, and these gradients can enable energy saving if fish can position themselves appropriately: drag forces can be counteracted by lift-based thrust generation. We obtained simultaneous kinematic and metabolic data on rainbow trout swimming both in free stream flow and in a velocity gradient generated by an airfoil within a respirometer. As water moves past the sharp edges of the airfoil, a steep velocity gradient is generated, and trout will voluntarily position themselves in this gradient region, orient their bodies at a small angle of attack, and greatly reduce body movements. We simultaneously and continuously measure how fish respond metabolically and kinematically when they volitionally move from the freestream to the region of sharp velocity gradient to quantify how variation in kinematics induced by the velocity gradient translates into metabolic variation.

Utilizing the IBAS respirometry system for measuring kinematic and metabolic data simultaneously and the dynamic metabolic rate measuring technique, we were able to observe how fish modulate their dynamic patterns of kinematics and metabolism when they respond to a sharp fluid velocity gradient (Fig. 4). When fish swim in free stream water, velocity is at 50% of their maximum sustained swimming speed, and fish have a dynamic metabolic rate at the level of ∼250 mg O_2_ kg^-1^ h^-1^. The peaks (∼290 mg O_2_ kg^-1^ h^-1^1) and the troughs (∼205 mg O_2_ kg^-1^ h^-1^) of the metabolic rate (Fig. 4a) are reflected in the magnitude of tail displacement (Fig. 4b) (25 to 75 percentile: 220-252 mg O_2_ kg^-1^ h^-1^; coefficient of variation: 8.3%) as fish swim actively against the flow. When fish move into the fluid velocity gradient and can maintain position with minimal body movement by using their body to generate lift and hence thrust, metabolic rate drops 76% to ∼60 mg O_2_ kg^-1^ h^-1^ (25 to 75 percentile: 60-69 mg O_2_ kg^-1^ h^-1^; coefficient of variation: 13.7%) (Fig. 4a). Correspondingly, tracking tail movement revealed that variation in the magnitude of tail displacement reduced substantially and the tail movements observed result from largely passive body movements resulting from incident flow (Fig. 4b) when fish are in the velocity gradient. Therefore, fish in the velocity gradient have a much smaller magnitude of tail displacement and lower metabolic rate than when fish are in the free stream.

**Figure 4.**
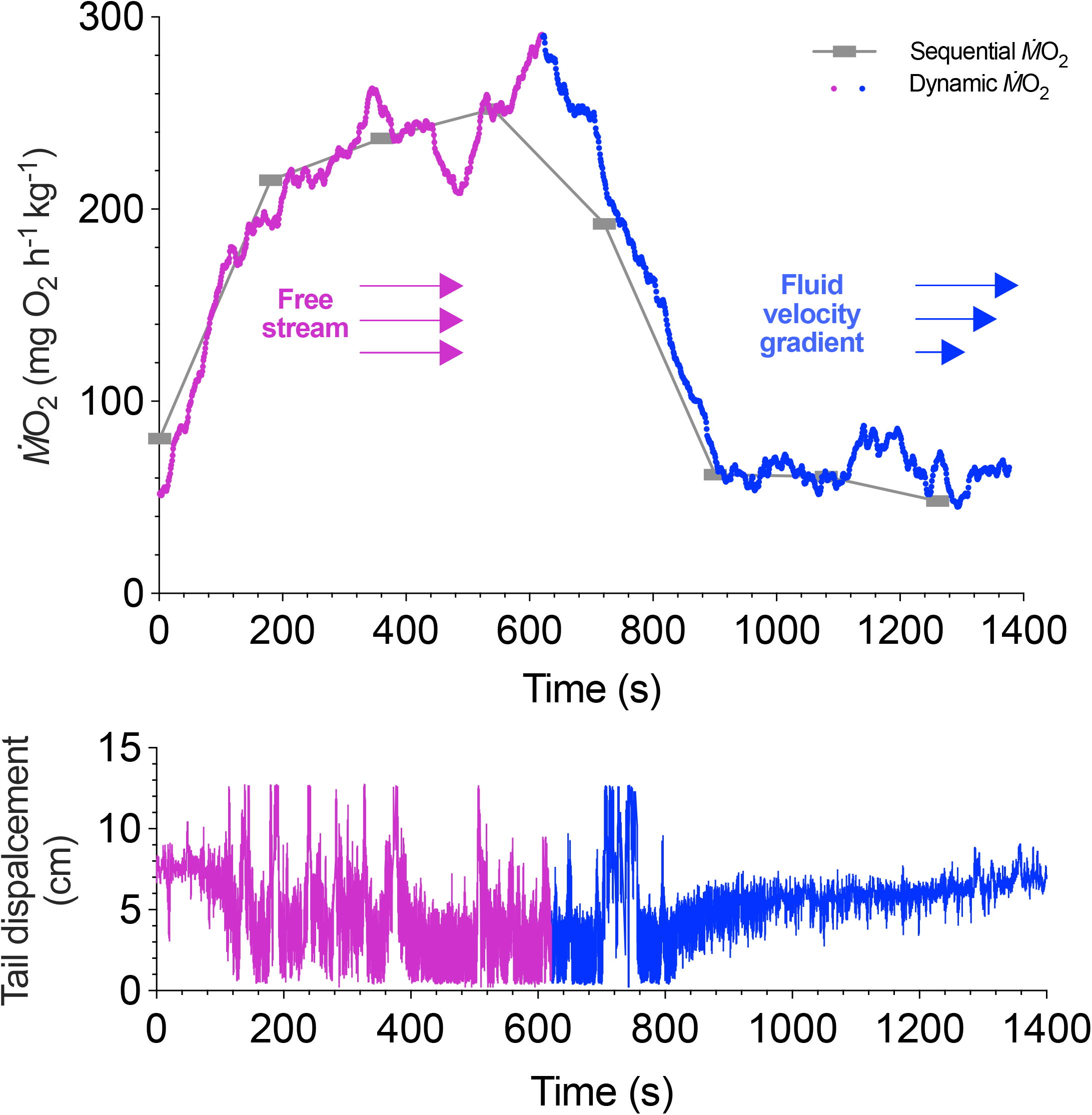
Temporal matching of measured dynamic aerobic metabolic rate with locomotion of a trout moving from the free stream and into a velocity gradient where the fish can maintain position with little metabolic cost. (a) Dynamic aerobic metabolic rate (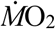, rate of oxygen uptake) of a brook trout (*Salvelinus fontinalis*) changes with the fluid conditions, from the free stream (50% of the sustained maximum swimming speed) (purple line) to regions of sharp fluid velocity gradient (blue line). The group of arrows is a graphic illustrations of the flow velocity gradient. The conventional method of measuring aerobic metabolic rate using a sequential algorithm is provided as a reference (grey bars). (b) Tail displacement amplitude within the test chamber temporally matched to measurement of aerobic metabolic rate shown in (a). Movement into the velocity gradient results in reduced amplitude and lower variation tail motion.

More broadly, fish in the natural environment encounter variations in the flow field, which may contribute to variations in their kinematics and energetics. For example, fish swimming in turbulent conditions encounter vortices across different scales [32]. Fish in schools also face non-uniform fluid conditions. As their neighbours and leaders undulate their bodies, they shed periodic vortices that the following fish may take advantage of, similar to the fish swimming behind a hydrofoil [38].

## 6. Conclusions

Animals in nature rarely show constant locomotion with low variation; instead, animals actively modulate their locomotion when biological and environmental conditions change. Other than the manoeuvrability needed for predator-prey interactions or collision avoidance, why do animals modulate their locomotion? We propose that modulated locomotion is coupled with dynamic changes in metabolism to regulate and reduce the overall metabolic cost of movement. By altering patterns of locomotion to take advantage of local fluid dynamic conditions and hence to achieve energy saving, animals will have more metabolic capacity available for other fitness-related functions. Variation in biomechanics and metabolism is coupled, and we believe that deeply integrating the analysis of biomechanics and bioenergetics by synchronized measurements will allow us to better understand the effects of observed variation in kinematics on energetic cost. We show that fish within a school actively modulate their locomotor dynamics, changing both frequency and amplitude, and we demonstrate that fish swimming in velocity gradients show dramatic energetic savings by taking advantage of fluid dynamic pressure gradients. Studies that synchronize and continuously measure energetics and kinematics provide a basis for understanding the ubiquitous variation that is observed in fish locomotion and reflect the near-constant dynamic adjustments that animals make to navigate a complex and constantly changing environment.

## Ethics

This work has ethical approval from Harvard Animal Care IACUC Committee (protocol number 20-03-3).

## Data accessibility

Data are archived in a repository for publication.

## Funding information

Y.Z. is supported by a Postdoctoral Fellowship of the Natural Sciences and Engineering Research Council of Canada (NSERC PDF-557785–2021), followed by a Banting Postdoctoral Fellowship (202309BPF-510048-BNE-295921) of NSERC & CIHR (Canadian Institutes of Health Research). Funding provided by the National Science Foundation grant 1830881 (GVL), the Office of Naval Research grants N00014-21-1-2661 (GVL), N00014-16-1-2515 (GVL), 00014-22-1-2616 (GVL).

## Acknowledgments

Many thanks to members of Lauder Laboratory for numerous discussions, to Prof. Tony Farrell for the work together for the discovery of dynamic metabolic rate, and to Cory Hahn for fish care.

## Author contributions

Y.Z., G.V.L. conceptualized the study. Y.Z. performed experiments and data analyses and wrote the original manuscript. D.R., H.K. contributed to data analyses and visualization. All authors provided manuscript edits and comments and approved the final version.

## Competing interests

Authors declare that they have no competing interests.

## Notes

### Competing Interest Statement

The authors have declared no competing interest.

